# Inversion characterization in *Timema* stick insects: from local PCA to pangenome approaches

**DOI:** 10.64898/2026.09.18.752789

**Authors:** Diana Tataru, Kira Vasquez-Kapit, Patrik Nosil, Zachariah Gompert

## Abstract

Structural variants (SVs) can strongly influence evolutionary processes by suppressing recombination to create jointly inherited regions and by altering gene expression and the 3D organization of DNA. Recent improvements in sequencing technologies have led to an influx of high-resolution genomic data, bolstering research on SVs. When detecting and characterizing SVs, researchers face trade-offs in data and methods concerning scale, resolution, and computational intensity. Here, we evaluate such trade-offs by quantifying how data type, analytical approach, and SV attributes affect the ability to detect and characterize SVs in *Timema cristinae* stick insects, with a focus on inversions. We specifically evaluate inversions characterized by (i) a population genomic approach based on patterns of local population structure inferred from genotyping-by-sequencing data, (ii) pairwise alignments of eight *de novo* genome assemblies and SV calling with SyRI, and (iii) a pangenome constructed from the alignment of the eight genome assemblies. Our results suggest that while broad genomic regions harboring SVs can be detected by any of the approaches, the approaches differ in how they characterize inversions, especially small or complex ones. The local population structure approach is exploratory and generally only detects the largest inversions. Pairwise comparative alignments identify the greatest number of inversions, but downstream analyses are influenced by identification of shared inversions across genome pairs. Pangenome approaches are scalable and produce network graphs that describe complex SVs but require a conceptual shift in visualization and interpretation. The collective results highlight limitations and benefits of different approaches in a burgeoning research field.

## Introduction

The field of comparative genomics and structural variant (SV) detection is growing quickly. Studies are continually identifying the large impacts of structural variants on critical evolutionary processes (Todesco, et al., 2020; Tigano, et al., 2021; Hamala, et al., 2021; Merot, et al., 2023; Kollar, et al., 2025; Stuart, et al., 2025; Gomez-Ramos, et al., 2026). SVs can cause recombination suppression over large genomic regions leading to co-adapted gene complexes, alter the 3D structure of DNA, and cause functional changes in the genome (Stuart, et al., 2025). However, SV detection and characterization have long posed significant challenges to researchers. With the increasing availability of many different types of genomic sequencing data and the development of new SV detection tools, the question remains of how genomic data types and structural variant detection methods affect SV identification. Specifically, are we missing or misrepresenting SVs when we choose a specific method and how does SV size, frequency, and complexity influence detection by different methods?

The distinction between detection and characterization is critical, as these different aspects answer different questions. Detection answers “is there structural variation here?” while characterization answers “what are the properties of the SV?”, including SV type, boundaries of individual SV elements, and delineating SV signals into constituent structural elements. Detection and characterization can occur on different scales. For example, pairwise whole-genome alignments might miss an SV due to limited number of genomes (some SVs not being present in the sequenced samples) but identify the exact breakpoints and components of a detected SV. Existing comparisons of SV detection among methods have confirmed bias in SV detection by data type, software, and an interaction of data type and software (Meng, et al., 2023; Merot, et al., 2023). Comparisons of both short-read (Cameron, Di Stefano, & Papenfuss, 2019) and long-read (Lucek, Gompert, & Nosil, 2019; Liu, Xie, & Li, 2024) SV callers found widely varying detection across SV callers. The resolution of empirical data needed to compare these SV characterization approaches across populations and data types is still mostly lacking in most biological systems, but the rapid generation of new sequencing data makes this comparison necessary and timely.

In this study, we use population level genotyping-by-sequencing data and multiple whole-genome assemblies from a system in which SVs have been shown to underlie adaptive traits. We describe differences among newly developed and widely-used SV detection methods, thereby highlighting advantages of each. We specifically focus on inversion detection methods using: (1) local population structure inferred from genotyping-by-sequencing (‘local PCA’ hereafter, where PCA refers to principal components analysis), pairwise whole-genome comparative alignments, and recently developed pangenome SV calling. As outlined in more detail below, all three of these methods have benefits and limitations relating to their cost, resolution, and relevance to population-wide or species-wide patterns.

Local PCAs detect outliers in windowed principal components analyses across the genome and characterize SVs as regions where a series of SNPs with similar patterns of population genetic structure occur (Li & Ralph, 2019; Huang, et al., 2020). While runs of PCA outliers do suggest limited recombination in a region, this approach only infers SVs from population structure, rather than identifying them directly. Thus, outlier runs require further confirmation through other methods to verify SV presence as opposed to recombination suppression for other reasons (e.g. proximity to centromere, meiotic drive). Moreover, outlier runs do not characterize SV nature, complexity, or size on their own. Nonetheless, local PCA approaches are a common way to evaluate SVs in populations using, but not restricted to, genotyping-by-sequencing (GBS hereafter) data (Li & Ralph, 2019; Huang, et al., 2020; Todesco, et al., 2020; Harringmeyer & Hoekstra, 2022; Petak, et al., 2026). GBS data include a range of reduced representation sequencing data types varying in enzyme digestion, adaptor ligation, barcoding, and size selection (e.g. GBS, RADseq, 2bRAD, ddRADseq), resulting in relatively affordable data which can be generated for a large number of samples (Andrews, Good, Miller, Luikart, & Hohenlohe, 2016). Thus, local PCAs can give a better idea of putative SV frequency in a population relative to approaches that use only a handful of genomes. However, past work in our system focused on a small subset of genome-wide variation and large inversions found that the local PCAs only detected about half of the inversions present in the comparative alignments (Gompert, et al., 2025). These discrepancies may be due to factors like haplotype frequency within population, or variability within population, size, or inversion age (e.g., young inversions may yet to have accumulated substantial genetic structure) leading to limited differentiation in population structure.

Comparative alignments refer to pairwise comparisons of synteny using independently derived whole-genome assemblies, producing detailed and easily viewable and interpretable pairwise outputs (e.g., ‘dot plots’), which can be analyzed with downstream SV callers and clustered across pairs. Whole-genome assemblies are expensive to generate relative to GBS and remain relatively limited in number, so comparative alignments can miss or misrepresent some of the population variation that local PCAs identify. However, the number of available assemblies is increasing quickly (Lewin, Richards, Aiden, & et al, 2022; Darwin Tree of Life Project Consortium, 2022). Due to their pairwise nature, comparative alignments also lack some of the scalability of pangenomes, which incorporate all genome assemblies into one graph. Tree based approaches in genome alignment can integrate multiple genomes (Armstrong, et al., 2020), and methods do exist to align multiple genomes to one reference without creating a pangenome (Minkin & Medvedev, 2020; Lovell, et al., 2022). These are more scalable than simple pairwise alignments but require genomes to not be too diverged (Minkin & Medvedev, 2020) or have conserved gene content (Lovell, et al., 2022). This is limiting considering one of the major draws of comparative alignments is that, unlike local PCA and some pangenome approaches, they can be done at many scales including within populations, between populations, or between species.

Pangenome graphs are networks of nodes, edges, and paths that describe variation in a collection of related whole-genome assemblies. Super pangenomes can compare genomes across species in a genus, but construction of super pangenomes balances a critical tradeoff between haplotype resolution in closely related genomes and genetic diversity across highly divergent species, where each can substantially increase computational complexity (Raza, et al., 2026). Pangenomes are the newest approach in SV detection and are promising for describing large-scale and complex SVs, but remain limited in their development (Hickey, et al., 2020). Pangenome graph construction can be computationally intensive, and for this reason graphs are often stored and analyzed chromosome-by-chromosome, making interchromosomal rearrangements more difficult to quantify. In pangenome graphs, SVs are defined as topological motifs called bubbles that are not annotated for SV type (Hickey, et al., 2024; Romain, et al., 2025). Nested variants appear as nested bubbles, defining a layout for complex SVs. Pangenomes also solve the issue of reference bias and can have better alignment than linear references (Secomandi, et al., 2025; Garrison, et al., 2018). Studies have found that SV genotyping is significantly improved when short read sequencing data is aligned to a pangenome rather than an individual reference (Hickey, et al., 2020), but the inability to create joint VCF (Variant Call Format) files across individuals means that classic population genomics approaches such as local PCA outlier detection are then challenging to implement. For this reason, we examine the pangenome and local PCA approaches separately, because although they will likely become complementary, they are currently most commonly used separately.

We focus on characterization and detection of inversions using these three approaches in *Timema cristinae* (stick insects) located on two nearby mountains. This system is well-suited for such an evaluation because the required data types exist (e.g., population-level GBS data plus replicated phased genome assemblies) and there is known structural variation. For example, complex structural variation on Chromosome 8 is associated with variation in cryptic coloration (Gompert, et al., 2025; Nosil, et al., 2024; Nosil, et al., 2018). Specifically, complex structural variation distinguishes striped versus unstriped morphs. These morphs bear a longitudinal dorsal white stripe and are adapted to the host-plant *Adenostoma fasciculatum,* versus are solid green and adapted to *Ceanothus spinosus*, respectively. On the two mountains we investigate in this study, Refugio and Hwy 154, both morphs occur in sympatry (e.g., often on the exact same plant individual, for example due to gene flow between populations on different hosts). In addition to the complex SVs (e.g., inverted translocations) on Chromosome 8 that associate with cryptic coloration, past work also revealed substantial SVs on other chromosomes that are not associated with coloration, setting the stage for the current evaluation of genome-wide SV detection and characterization using different approaches (Gompert, et al., 2025; Nosil, et al., 2024; Nosil, et al., 2018). We focus here on inversions as a logical starting point because they exist in many systems (Tigano, et al., 2021; Hirabayashi & Owens, 2023; Zhou, et al., 2023; Gozashti, Harringmeyer, & Hoekstra, 2025) and they substantially reorganize chromosome structure and suppress recombination such that they are particularly relevant to studies of adaptation and speciation.

In summary, we present a comparison of three methods of inversion detection and characterization with different types of genomic sequencing data, to better inform a quickly developing field. We also describe how inversion size, density, and complexity interact with the ability to detect and characterize inversions. We chose the specific methods for each approach based on what is currently common in the literature (local PCA and comparative alignment) or what represents a likely major avenue for future studies (pangenome). Although our comparison does not fully encapsulate all the possible methods for SV detection and characterization, it nevertheless serves as a logical starting point to inform researchers moving forward.

## Methods

### Sequencing Data

We analyzed *T. cristinae* DNA sequencing data from two mountains outside of Santa Barbara, CA, USA; Refugio and Hwy 154. Hwy 154 is represented by a large host-plant patch called FHA (latitude 34.5324◦N, longitude 119.8473◦W) while Refugio is represented by a series of closely-connected localities described in detail in Gompert et al. 2025. We analyzed whole-genome-sequencing (WGS) of eight *de novo* haploid genome assemblies derived from four individuals (four genomes from each mountain, two per host plant) generated as described in (Gompert, et al., 2025; NCBI PRJNA1208027 to PRJNA1208034). Briefly, haplotype-phased chromosome-level genomes were constructed by Cantata Bio using a combination of PacBio and Illumina reads from Omni-C genomic libraries. We also analyzed GBS data from the two mountains generated as described in past work (Gompert, et al., 2025; Comeault, et al., 2015). This consisted of single-end 150 base-pair data of 238 *T. cristinae* insects from Refugio and 602 *T. cristinae* insects from Hwy 154.

### Local Population Structure Analysis

To detect series of outliers of within-population variation (inferred inversions) in the GBS data, we used a local PCA approach. We first conducted a variant calling workflow separately for each mountain. We aligned GBS data to the GSH2 (Hwy 154 Stripe haplotype 2) reference genome from Gompert et al. 2025 with bwa aln/samse version 0.7.19 (Li & Durbin, 2009), sorted and indexed with samtools version 1.16 (Li, et al., 2009; Danecek, et al., 2021), and called variants with bcftools mpileup/call version 1.16 (Danecek, et al., 2021). We filtered variants with custom scripts from (Gompert, et al., 2025), with thresholds described in SI, and then used an empirical Bayesian approach to estimate expected genotype frequencies given allele frequencies and random mating expectations, as described in the Supplementary Information (SI) and a previous publication (Soria-Carrasco, et al., 2014). After filtering, this resulted in 146,886 SNPs from 236 individuals for Refugio and 329,268 SNPs from 565 individuals for Hwy 154.

We analyzed population structure across the genome and identified outliers (i.e., sequential regions with accentuated structure) using the R package lostruct (Li & Ralph, 2019), following filtering slightly altered from (Huang, et al., 2020; https://github.com/dianatataru/Timema-SV-Methods/blob/main/local_pca/localpca_manyMDSaxes_withfiltering.R). All R analyses were conducted in R version 4.5.1 (Team, 2025). We standardized data, ran a principal component analysis (PCA) on each non-overlapping window of 100 SNPs to identify structure within windows, and calculated Euclidean distance between windows for the first two principal component axes. We then mapped distances between windows for PC1 and PC2 with a multidimensional scaling (MDS) analysis in 40 dimensions, and defined outlier windows as those with absolute values greater than three standard deviations from the mean across all windows for each MDS dimension (Figure 1A).

**Figure 1.**
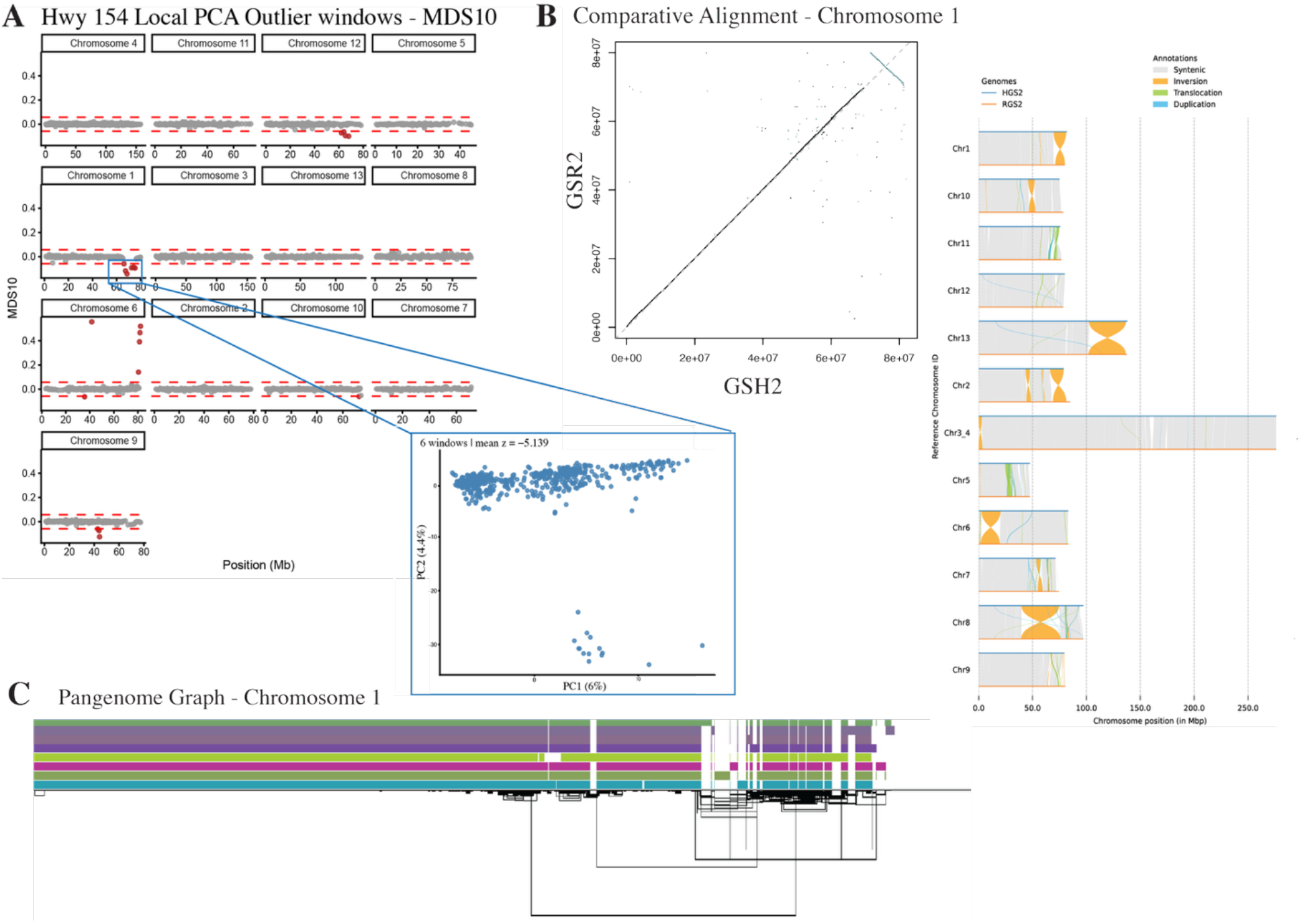
Conceptual figure of three structural variant calling approaches. (A) Local PCA outliers for genotyping-by-sequencing data from one of the two mountains (Hwy154) on MDS 10 (left) and the corresponding PCA for that outlier region, showing three clusters (right). (B) Pairwise comparative alignment between two *de novo* whole-genome assemblies (GSH2 & GSR2) for Chromosome 1 (left), with structural variant calls across chromosomes from SyRI annotated for inversions, translocations, and duplications (right). See Supplementary Figure 4 for other six pairwise comparisons. (C) Example of pangenome graph for Chromosome 1 constructed from the eight genome assemblies visualized in ODGI viz (Optimized Dynamic Genome Graph Implementation visualization; https://github.com/vgteam/odgi). Each bar represents a path (genome) and lines (links) represent graph topology. See Supplementary Figure 5 for all chromosomes.

We clustered windows within and between populations as described in SI and ran a permutation to test if the longest run of outlier regions on an MDS axis is longer than expected by random chance under the null hypothesis of no autocorrelation along the genome (n = 1000, p threshold = 0.01), only keeping significant regions. Lastly, we applied a minimum filter to retain only sets of five or more consecutive outlier windows. Note that the genomic regions we detect with this method are putative inversions, meaning we know they are probably regions of reduced recombination and possibly SVs, but we are less able to confirm the nature, complexity or causes of structure with this method. Additionally, SV size estimates from this approach are necessarily affected by window size and minimum window threshold.

### Whole-Genome Comparative Alignment

To detect and characterize inversions using a pairwise comparative alignment approach, we used the program Progressive Cactus (version 2.7.2; Armstrong, et al., 2020), comparing seven reference-quality genome assemblies pairwise to the GSH2 genome assembly, the same genome used as the reference for the local PCA analyses. We made pairwise dot plots with Progressive Cactus (Armstrong, et al., 2020) for visualization. We then aligned the pairwise genome assemblies with minimap2 (version 2.22; Li, H., 2018; Li, H., 2021) and used the program SyRI version 1.8.2 to call pairwise SV summaries (Figure 1B; Goel, Sun, Jiao, & Schneeberger, 2019). SyRI identifies the longest syntenic path between a pair of genomes and then classifies aligned non-syntenic regions as either an inversion, a translocation, a duplication, or a combination of the three (i.e., a complex SV).

To cluster across the pairwise SyRI output, we used a custom R script (https://github.com/dianatataru/Timema-SV-Methods/blob/main/inversion.R) to combine inversions across pairwise comparisons if all inversions in the cluster are at least 2/3 the size of the largest inversion in the cluster, and 80% overlapping in reference position with the largest inversion in the cluster, starting with the largest inversions. We calculate start and end positions as minimum start position across a cluster and maximum end position across cluster, thus generating an approximate maximum size estimate for each SV. We focus on inversions greater than 50 base pairs to have an equivalent threshold as the pangenome analysis (see SI for information on other SV types, e.g. translocations, duplications).

### Pangenome Analysis

Lastly, we used a pangenome approach to detect and characterize inversions, created using the same eight haplotype phased whole-genome assemblies as the comparative alignment, in Minigraph-Cactus version 3.0.1 (Hickey, et al., 2024). Minigraph-Cactus iteratively conducts pairwise alignments to an assigned reference genome using SVs greater than 50 base pairs in graph construction, calls SVs from all input assemblies, and creates a pangenome graph that individual genome paths are mapped back onto (Hickey, et al., 2024). For computational efficiency, we generated the pangenome graph chromosome-by-chromosome (Figure 1C). We built pangenome graphs with all four Hwy 154 genomes assemblies and evaluated pangenome graphs using the program Gretl (Variation GRaph Evaluation TooLkit version 0.1.1; Vorbrugg, et al., 2025). We used the reference genome with the longest pangenome graph (Hickey, et al., 2024). We visualized the pangenome using VG (version 1.67.0; Garrison, et al., 2018), ODGI (Optimized Dynamic Genome Graph Implementation version 0.9.4; https://github.com/vgteam/odgi), and Sequence Tube Map (Figure 2; Beyer, et al., 2019).

**Figure 2.**
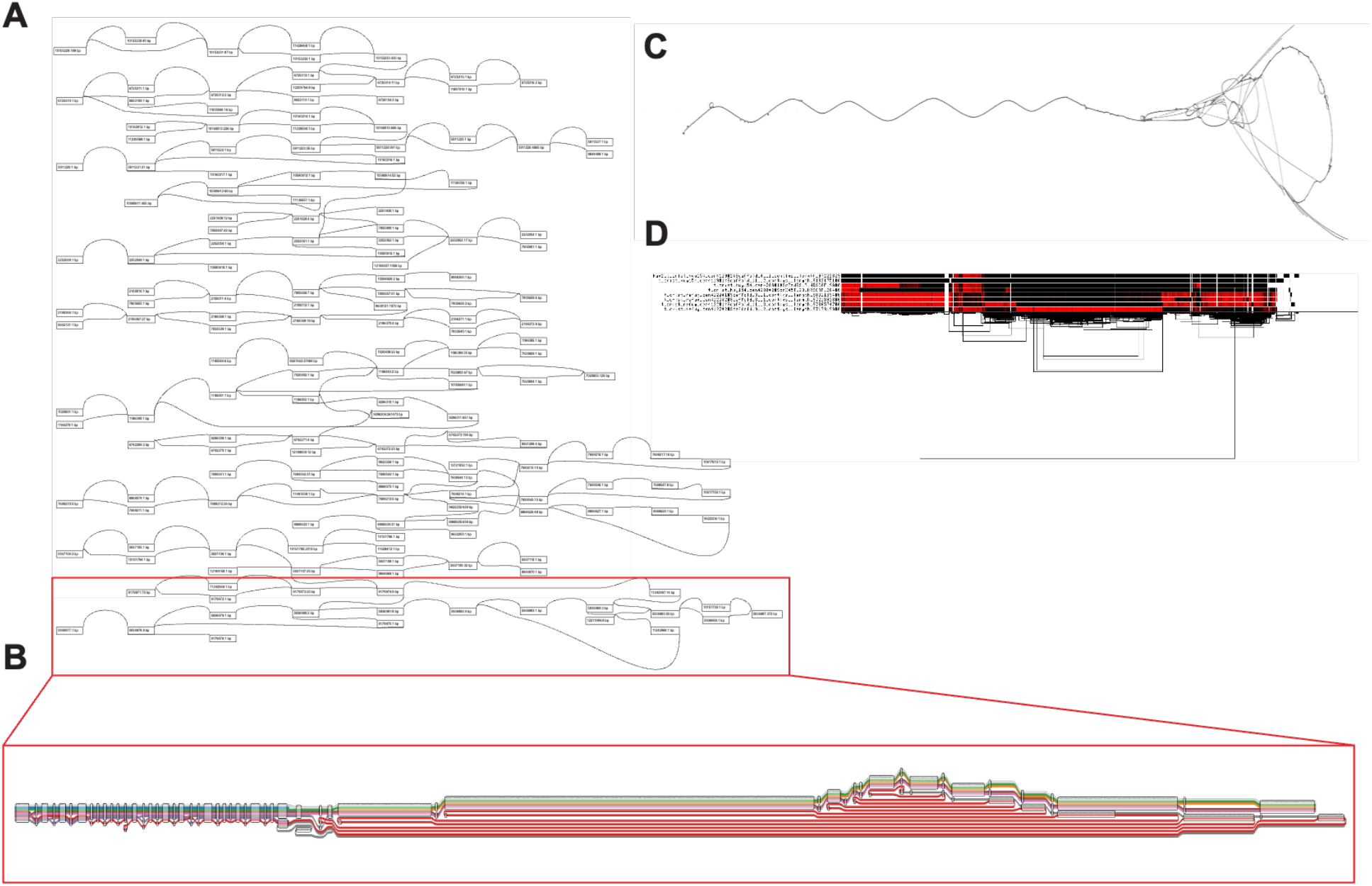
Visualization of Chromsome 8 of the pangenome graph. (A) Visualization of the nodes in VG viz (Variation Graph visualization; Garrison, et al., 2018) identified as inversion bounds using PanTree for Chromosome 8, with length of node in base pairs next to node number. Lines between nodes indicate edges, and visualization is expanded by three levels (-c 3) (B) Sequence Tube Map (Beyer, et al., 2019) visualization of the final inversion (nodes 9179861-9179887) pictured in (A), where each colored bar is a genome path. (C) Visualization of the entire Chromosome 8 with ODGI draw (Optimized Dynamic Genome Graph Implementation draw; https://github.com/vgteam/odgi), where lines indicate paths and loops indicate nested structural variation. (D) Visualization of the entire Chromosome 8 with ODGI viz (Optimized Dynamic Genome Graph Implementation visualization; https://github.com/vgteam/odgi), sorted by path-guided stochastic gradient descent, where each bar represents a path (genome), colored by whether it is inverted relative to the reference (red), and lines beneath the paths represent links, or graph topology.

Once the pangenome graph was built and evaluated, we characterized SVs in the pangenome using the program PanTree (version 0.3.0; Nowbandegani, et al., 2025). We also tried a second inversion-specific pangenome SV caller, INVPG-annot (version 1.1.0; Romain, et al., 2025), but found PanTree to be more robust (see SI for comparisons). PanTree calls variants in a pangenome by creating a reference tree with all of the nodes in the graph, but a subset of edges formed by traversing the graph while prioritizing the linear reference, defined as reference edges. Starting with the outer reference tree structure, PanTree defines variant edges relative to the reference tree into five non-overlapping categories: insertions, deletions, replacements, duplications, and inversions. Translocations are not specifically denoted in this method but could appear as a combination of a deletion and insertion. We subset the output VCF file from PanTree to inversions and extracted information using bcftools version 1.16 (Danecek, et al., 2021). Then, we used VG version 1.67 (Garrison, et al., 2018) to project the pangenome node bounds for inversions back into the linear genome coordinate spaces. This linearization of the pangenome greatly reduced the complexity of the graph but is necessary for comparing findings to the other two approaches. We estimated size of the inversion by taking the larger of the reference vs. alternate path lengths as described in SI. Again, here we estimate SV size as the largest of the two.

### Comparing Structural Variant Detection and Characterization (i.e., overlap among approaches)

To compare structural variant detection and characterization across our three methods, we calculated overlap in inversions across methods using three different approaches in a custom R script (https://github.com/dianatataru/Timema-SV-Methods/blob/main/inversion.R): 1) inversions are considered overlapping across methods when the smallest inversion is at least 2/3 the size and at least 80% positionally overlapping across the genome with the largest inversion in the cluster (i.e., group of inversions identified as overlapping across methods); this follows the within-method clustering criteria used for local PCA and comparative alignment and is the most conservative; 2) any overlap in positions across the genomes, which further groups inversions within methods because this criterion is less conservative than the threshold originally used; and 3) overlap as a proportion of the total genome, which does not differentiate between separate inversions that may be overlapping but structurally differentiated. While the third overlap method addresses detection, the first two methods handle differences in characterization, especially to the extent that characterization involves delineating SV elements and breakpoints.

We visualized overlap across methods using Venn diagrams in R package ggvenn (version 0.1.19; Yan, 2025). We modeled differences in inversion size between methods and number of methods detecting individual inversions with non-parametric Kruskal-Wallis tests and post hoc Dunn tests with Bonferroni correction in the R package dunn.test (version 1.4.1; Dinno, 2026). To test whether local inversion density affects detection across methods, we fit a logistic regression (binomial GLM) predicting detection as a function of inversion density (number of neighboring inversions within 100 kilobases) and method, as well as their interaction. We also fit a logistic regression with the following variables and their interaction with method; distance to nearest inversion (AIC = 1,604.2, residual deviance = 1,592.2) and number of inverted bases within 100 kb (AIC=1,607.7, residual deviance = 1,595.7), but number of neighboring inversions had the best model fit (AIC=1,485.1, residual deviance = 1,473.1).

## Results

### Local Population Structure

We used population level GBS data to identify outlier blocks of genetic variation (i.e., clusters in PCA space) across the genome within the two mountains, Refugio and Hwy154. After filtering variants, we ended up with 236 *T. cristinae* insects from Refugio and 565 *T. cristinae* insects from Hwy 154 for local population structure analyses. Before merging overlapping inversions within mountain (Hwy 154 or Refugio), we had 24 inversions for Refugio and 49 inversions for Hwy 154. In this count, inversions in the same genomic location but with different directionality (positive or negative z score) on the same MDS axis are considered different SVs. After merging within mountains (across negative/positive directions and MDS axes by any overlap in genomic position), we had 13 Refugio inversions (Supplementary Figure 1) and 23 Hwy 154 inversions (Supplementary Figure 2). After merging overlapping inversions between mountains with at least 2/3 size and 80% position overlap, we identified four inversions as shared between mountains (Supplementary Figure 3). Thus, 4 of 36 total inversion calls (11%) were shared between mountains.

Overall, we detected 32 putative inversions across the 13 chromosomes and two mountains (Figure 3A) with a mean inversion size of 11,162,106 base pairs (range = 1,947,016 – 52,204,874 bp). The proportion of the genome covered by these putative inversions (i.e., local PCA outlier windows) was 25.78%.

**Figure 3.**
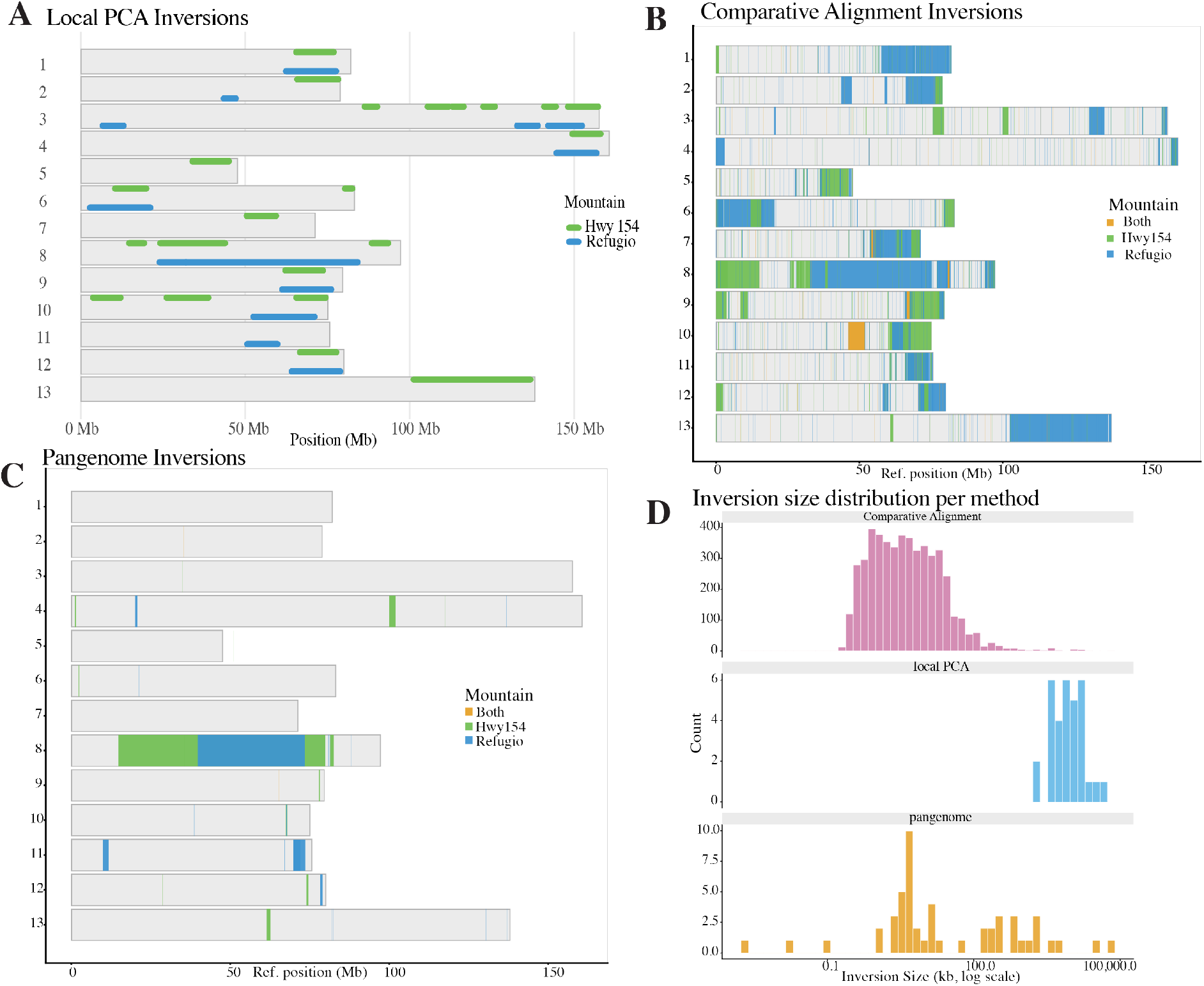
Summary of inversions found across the three methods in the reference genome (GSH2) coordinate space, where inversions are colored by mountain (Hwy 154 = green, Refugio = blue, both = orange). (A) Local PCA outliers called with lostruct (Li & Ralph, 2019), (B) comparative alignment inversions called using the program SyRI (Goel, et al, 2019), (C) pangenome inversions called using the program PanTree (Nowbandegani, et al., 2025), and (D) inversion size distribution with size represented as log scale kilobases and count. Note the different y axis for each method: comparative alignment (pink; 0-400), local PCA (blue; 0-6), and pangenome (orange; 0-10).

### Whole-Genome Comparative Alignment

Using comparative alignments to detect and characterize SVs across genome assemblies, we found a higher number of inversions relative to the other two methods. The number of SVs detected were sensitive to clustering methods (i.e., how inversions called between pairs overlapped with those called between other pairs), which were necessary given the pairwise nature of the analyses. Before clustering and filtering we detected 7,020 inversions (Supplementary Figure 4). After clustering by 80% position and 2/3 size overlap and filtering to a minimum size of 50 base pairs to match pangenome filtering, we detected 4,904 inversions across the seven pairwise alignments (Figure 3B), with mean inversions size of 74,070 base pairs (range = 201 – 42,064,975 bp). Of these 4,904 inversions, 1,619 (33%) contain inverted translocations, 2,070 (42%) contain inverted duplications, 300 (6%) contain both inverted duplications and translocations, and 902 (18%) are simple inversions. This suggests that most of the inversions (82%) are complex. For this method, inversions covered almost a quarter 23.27% of the total genome. This is a similar percentage of the genome to the local PCA method, despite the larger number of SVs called in the comparative alignment. This discrepancy in number of SVs relates to inversion size, and the higher likelihood of calling smaller inversions in the comparative alignment relative to the other two methods.

### Pangenome

Using a pangenome graph and the program PanTree to detect and characterize SVs, we identified 51 inversions across the 13 chromosomes (Figure 3C), which covered 6.4% of the genome. This reported coverage is substantially smaller than the other two methods. Given our projection back on the linear reference genome, mean inversion size was 2,357,001 base pairs (range = 2 – 65,057,107 bp), with 49 of 51 inversions greater than 100 base pairs in size. Note the minimum range is less than the 50 bp threshold because the linearized size metric does not calculate size exactly. The total number of inverted base pairs from the PanTree output is 120,207,099 base pairs, calculated from the size metric we determined by choosing the longer of the linearized reference and alternate paths. The Gretl analysis of the full pangenome identified total inverted nodes for each haplotype genome path to be between 389,515,176 – 582,896,541 base pairs (Table 1). While the inverted nodes identified by Gretl are raw topology features and not variant calls like those from PanTree, the discrepancy between these numbers suggests that the PanTree inversions projected back into linear reference space underestimates the size of inverted regions across genomes. This helps explain the relatively small proportion of the genome covered by inversions relative to the other methods.

**Table 1.** Output of Gretl (Vorbrugg, et al., 2025) for each genome path within the pangenome graph, where GSH2 is the reference.

| **Genome** | **Haplo-**<br>**type** | **Sequence**<br>**length (bp)** | **# Nodes** | **# Edges** | **Inverted**<br>**nodes (bp)** | **PropGe-**<br>**nomeIn-**<br>**verted** |
| --- | --- | --- | --- | --- | --- | --- |
| hwy154_stripe | 1 | 1220429573 | 80552526 | 80552513 | 777165 | 0.0006 |
| hwy154_stripe | 2 | 1226560494 | 81287868 | 81287855 | 454245474 | 0.3703 |
| hwy154_green | 1 | 1204896739 | 80373753 | 80373740 | 462513309 | 0.3839 |
| hwy154_green | 2 | 1215314917 | 81051730 | 81051717 | 518414847 | 0.4266 |
| refug_green | 1 | 1239970858 | 73079069 | 73079057 | 452537233 | 0.3649 |
| refug_green | 2 | 1235469353 | 73266674 | 73266662 | 389515176 | 0.3153 |
| refug_stripe | 1 | 1227621598 | 80118808 | 80118795 | 582896541 | 0.4748 |
| refug_stripe | 2 | 918956906 | 62613904 | 62613893 | 566335954 | 0.6163 |

### Overlap among approaches: general trends

When comparing inversions across methods, we found that the extent to which inversion detection and characterization overlapped among methods varied based on how overlap was defined. For the most conservative definition of overlap (80% position and 2/3 size), there were no inversions shared by all three methods, one shared inversion between the local PCA and pangenome, five shared inversions between the local PCA and the comparative alignment, and twelve shared inversions between the pangenome and the comparative alignment (Figure 4A). This was no different than would be expected by random chance for inversions shared among all three methods, (Z-score = –0.56, p = 0.58) and between the local PCA and the pangenome (Z-score = 1.22, p = 0.11). We found fewer shared inversions than expected by random chance between the local PCA and the comparative alignment (Z-score = –5.26, p = 1.47e-07), and the pangenome and the comparative alignment (Z-score = –4.75, p = 2e-06). In summary, the most stringent thresholds for overlap resulted in very few overlapping inversions among methods.

**Figure 4.**
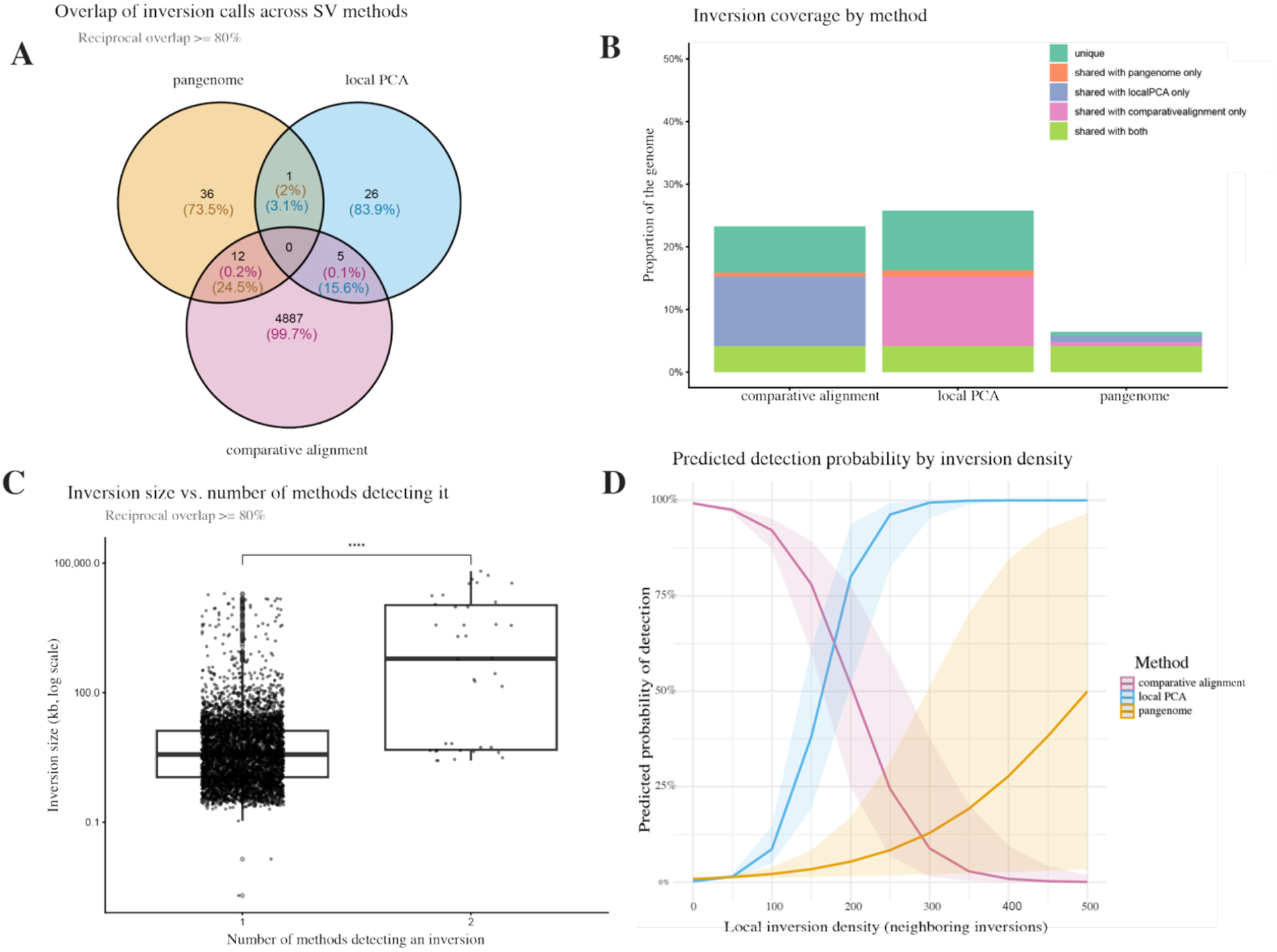
Overlap of inversions across methods. (A) Venn diagrams of shared inversion clusters between methods where clusters are >80% position overlapping and at least 2/3 the size of the largest inversion in the cluster. Percentage is colored by corresponding methods where pink is comparative alignment, blue is local PCA, and orange is pangenome. (B) Proportion of the genome covered by inversions for each method. Bar plots are stacked and colored by whether that inversion location is shared across methods. (C) Inversion size distribution by number of shared methods detecting every identified inversion. Pairwise significance is denoted where **** indicates p < 0.0001. (D) Binomial GLM predicted detection probability by inversion density in 100kb windows and method. The lines represent the predicted values from the model for each method and the shaded regions represent the 95% confidence intervals around predicted values.

For any overlap in position qualifying inversions as the same across methods, we found ten inversions shared among all three methods (Supplementary Figure 6). This is more than the first definition of overlap which found none, and significantly more than null expectations (Z-score = 16.45, p = 8.3e-61). There were no inversions shared between the local PCA and the pangenome, a pattern not significantly differing from the null (Z-score = –0.59, p = 0.56). There were seventeen inversions shared between the local PCA and the comparative alignment, significantly less than expected by the null (Z-score = –2.07, p = 0.039). The pangenome and the comparative alignment shared sixteen inversions, also significantly less than the null expectation (Z-score = –3.91, p = 9.35e-05). For this definition of any overlap, all except one of the local PCA inversions had overlap with the other two methods, almost 2/3 of the pangenome inversions had any overlap with the two methods, but still only about 1.5% of the comparative alignment inversions had any overlap with the other two methods. This suggests that inversions called by the local PCA and pangenome were largely shared between methods, and the lack of significance relative to the null expectations is largely driven by the large number of inversions detected by the comparative alignment.

For our last metric of overlap in inversions across methods, shared inverted bases by genome position, we found that 4.13% of the total inverted bases across the genome were shared across all three methods, significantly greater than the null expectation of 0.38% (Z-score = 21,207, p < 0.0001; Figure 4B). The local PCA estimated the largest proportion of genome as inversion (25.78%), closely followed by the comparative alignment (23.27%; Figure 4B). Between these two methods, the local PCA and the comparative alignment, 11.11% of the total genome was shared inverted bases, compared to null expectation of 6% (Z-score = 7,547, p < 0.0001; Figure 4B). For each of the two methods compared to the pangenome separately, we found a non-significant trend towards less than the null expectation (local PCA-pangenome= 1%, null = 1.64%; comparative alignment-pangenome = 0.62%, null =1.5%).

### Overlap among approaches: inversion size and density

We found significant differences in inversion size detected across methods (Figure 2D; Kruskal-Wallis chi-squared = 115.18, df = 2, p < 2.2e-16). Post-hoc pairwise comparisons using Dunn’s test with a Bonferroni adjustment showed that putative inversions inferred by local PCA were significantly larger than both the comparative alignment (Z-score = 9.74, p < 2.2e-16) and pangenome inversions (Z-score = 4.81, p < 2.2e-16). Inversions detected by the pangenome were significantly larger than the comparative alignment inversions (Z-score = –4.58, p < 2.2e-16). We also found significant differences in inversion size between being detected by one or two methods when overlap is quantified as 80% position and 2/3 size (Figure 3C; Wilcoxon rank sum W = 33,482, p-value = 7.475e-12) where inversions detected by two methods are larger (median= 611,953, inter-quartile range (IQR)=10,695,328, n=38) than those detected by one method (median= 3,697, IQR=11,972, n=4,949), with a small effect size (r = 0.0970).

Methods had variable inversion detection relative to inversion density in 100 kb windows (Figure 4D). Comparative alignment inversion detection slightly decreased with inversion density (β = −0.024, SE = 0.003, p < 0.001), while local PCA inversion detection significantly increased with increasing inversion density relative to comparative alignment (interaction β = 0.061, SE = 0.005, p < 0.001). Pangenome-based inversion detection was slightly increased with inversion density relative to the comparative alignment (interaction β = 0.033, SE = 0.005, p < 0.001).

### A note on previously a detected SV associated with cryptic coloration

All three methods detected a large SV in approximately the same region as the inverted translocations on Chromosome 8 that were previously shown to be associated with color-pattern variation in Gompert et al., 2025 (Supplementary Figure 7A). This region was estimated across four of the genomes by Gompert et al. 2025 to vary between 14.5 Mbp and 43.5 Mbp in size. After clustering within methods, the local PCA estimated the inferred inversion to be around 42 Mbp and the comparative alignment estimated the region to be around 52 Mbp. Gompert et al., 2025 did not cluster similar inversions across pairwise comparisons, which is likely why our comparative alignment estimate is larger. Our pangenome approach estimated a similar range as other methods for the large chromosome 8 inversion size (33.6 Mbp-65 Mbp).

### Overlap among approaches: summary

In summary, all three methods detected shared inversions, particularly larger ones, but the extent of this overlap varied greatly. Different methods detected and characterized inversions of different sizes and densities. The local PCA characterized larger and denser putative inversions relative to the other two methods. The comparative alignment characterized smaller and less dense inversions relative to the other two methods. Finally, the pangenome characterized inversions of intermediate size and density, but had a significantly lower proportion of the genome covered by inversions relative to the other two methods.

## Discussion

### Distinguishing between detection and characterization

Our study found that while the main regions harboring inversions were often detected by multiple methods, the breakpoints and structure of the inversions were largely different across methods. Inversion size played a significant role in whether inversions were detected across methods. Detection of an inversion by multiple methods was more likely for larger than for smaller inversions. A recent study with simulated data that quantified inversion detection for a pangenome versus comparative alignment also found that detection varied across inversion size and method, with decreased detection for small inversions for the comparative alignment and both small and large inversions for the pangenome (Romain, et al., 2025). This study also found decreased detection with increased inversion density (Romain, et al., 2025). Our comparative alignment inversion detection also decreased with density, but not the local PCA, for which there was higher detection for higher density regions. This overlap in patterns between simulated data and our empirical data is especially compelling and highlights the importance of SV characteristics in detection by different methods.

Methods also varied by how well they were able to characterize complex versus simple inversions, where a complex inversion consists of multiple types of combined SVs. Characterization is important because the community is increasingly finding that identified inversions are more complex than previously thought (Kollar, et al., 2025; Gompert, et al., 2025). We are not able to characterize complex SVs in the local PCA approach, which groups putative inversions in large blocks that the other two methods break up and doesn’t distinguish inversions from other mechanisms of suppressed recombination. On the other hand, the comparative alignment SVs was much more fragmented relative to the other two methods, resulting in far more reported inversions than the other two methods. These patterns across methods are confirmed in the density analysis. The comparative alignment classified different types of inversions and characterized only a minority (18%) of the inversions as “simple” (not also translocated or duplicated), something that is missed from the local PCA approach. The pangenome captured this complexity through different pangenome paths, but these paths are more challenging to clearly delineate as inversions with specific, linear breakpoints, and were not as obvious in our analysis.

### Inversions cover a large proportion of the genome

Two of our approaches (local PCA and comparative alignment) estimated that around one quarter of the genome contained inversions. Keeping in mind that this calculation was performed across multiple populations of a species, this number is similar to calculations in a recent rice pangenome (29%; Zhou, et al., 2023), deer mouse genome (30%; Gozashti, et al., 2025), and in Atlantic Silverside fish (23%; Tigano, et al., 2021). An analysis of paired plant genomes in the same genera used SyRI to identify inversions and found the total genome length in inversions to be between 1.3% (*Eucalyptus*) and 37.4% (*Fragaria*; Hirabayashi & Owens, 2023). Of course, different inversion calling methods will affect the amount of the genome reported to be covered by inversions, as seen by our results, so these comparisons should be interpreted cautiously. However, this suggests that our calculations fall within the range of other biological systems.

Interestingly the pangenome inversion estimates reported roughly a quarter the amount of the genome covered by inversions relative to other methods (6.4%). This number is drastically different from the proportion of inverted nodes in each genome path in the pangenome estimated by Gretl, which is closer to 30-60% of the genome (Table 1). Rather than representing an error, this vast difference between those two metrics represents the complexity of how one defines an inversion. What gets called an inversion is subjective to size cutoffs, clustering across samples, directionality relative to a reference, and frequency. Simply detecting inverted regions relative to a reference, which is what Gretl does, does not characterize an inversion. Future studies characterizing inversions should be explicit about how and why they define inversions as they do.

### Future Directions for Pangenome Graphs

Pangenome graph analysis, visualization, and interpretation is a quickly developing field (Bao & Weigel, 2025). Due to pangenome graphs’ abilities to conceptualize complex variation across whole-genome assemblies and species, we anticipate methods moving in this direction. While SV detection is established in pangenome graphs through bubble detection, annotation of SV type is still severely limited. We use a recently-developed program for annotation, PanTree, with a robust theoretical framework which accounts for nestedness (Nowbandegani, et al., 2025), but find that it still underestimates inversions relative to other methods in the way we have annotated them. This limitation in SV annotation is not unique to pangenomes; SVs can often be differentially characterized by different callers. For example, in our system, only the comparative alignment found inversions, notably inverted translocations, on the end of Chromosome 8 (Supplementary Figure 7A). It is possible that the complex nature of this SV is why it was not detected by the local PCA and pangenome approaches. These are ecologically relevant distinctions to make. In this system, this inverted translocation on the end of chromosome 8 was aligned with genomic association peaks for color morphology, highlighting the important ecological role of some of these SVs (Nosil, et al., 2024).

Annotation of complex SVs is not the only limit in pangenome development. Linear extrapolations also limit pangenome interpretation. In our overlap analysis, which projects the pangenome nodes back onto the linear reference, we do not capture all of the inverted regions in the pangenome graph. The linear projection suggests there are fewer and smaller inversions, but the reality is the complex matrix of the pangenome graph is best conceptualized as such, a complex matrix (Figure 2). We flattened our output to compare to the other two linear methods, which also limited overlap. This is not an oversight on our part, as identifying methods to compare pangenomes to existing data in the context of linear references will remain relevant, especially for visualization and downstream analyses, and should be further developed.

One application of the pangenome graph that we did not explore, to keep analyses relatively separate for comparison, is SV calling with GBS data from the pangenome (Hickey, et al., 2020). Once SVs have been identified in the pangenome, this approach allows for identification of population level variation of SVs with more abundant and lower coverage data. While it faces some of the limitations we have already discussed with SV annotation and visualization, once researchers have shifted to a conceptual framework of a graph matrix, this a major advantage of pangenomes and a likely future direction for the field.

### Conclusions

When determining how to define, detect, and characterize SVs, the methods researchers choose will largely depend on the question they are asking. If simply concerned with blocks of limited recombination (caused by putative SVs) within a population or maintenance of balanced polymorphisms, local PCA, which chooses the largest bounds, might be the most suitable method. But if one is more curious about specific breakpoints, or the pathway of evolution (comparing two distantly related species), comparative alignments might be more appropriate. Lastly, if the goal is to develop a scalable system to be used for many questions, or across less closely related organisms, a pangenome may be most appropriate, although there are limitations to the extent of pangenome software development. Like with many natural phenomena, as we get closer to describing the actual complexity of a system, we get farther from being able to simply categorize the components of that system. Rather than shying away from this newfound complexity, the field can hope to use it to better characterize SVs. In turn, this may lead to more detailed and comprehensive understanding of SVs and their role in evolution.

## Acknowledgements

Research was supported by funding from the National Institutes of Health (NIH MIRA award, FAIN R35GM158189, to Z.G; NIH Training Grant, T32GM132057, to K.V.) and the French Laboratory of Excellence project “TULIP” (ANR-10-LABX-41) funded by a government grant as part of the France 2030 program, as a Senior Package to PN. Its contents are solely the responsibility of the authors and do not necessarily represent the official views of the National Institutes of Health. Thank you to the Center for High Performance Computing at the University of Utah for support and computational resources.

## Data Availability

The eight reference genomes described in this study are available from the National Center for Biotechnology Information (NCBI) (BioProject accessions PRJNA1208027 to PRJNA1208034). Genotyping-by-sequencing data described in this study are also available from NCBI (NCBI PRJNA1211795, PRJNA243533, PRJNA1010130, and PRJNA284835). All analysis scripts are available from GitHub (https://github.com/dianatataru/Timema-SV-Methods) with final versions archived on Dryad.

