## Supplementary material for "Inversion characterization in *Timema* stick insects: from local PCA to pangenome approaches": SI

### Supplementary Materials

#### Genotyping-by-sequencing Data Variant Calling and Filtering

We aligned samples from each mountain separately to the GSH2 reference genome (NCBI PRJNA1208029) with bwa aln/samse version 0.7.19 (Li & Durbin, 2009), with maximum four total mismatches and maximum two mismatches in a 20 base pair seed. We trimmed bases with quality scores less than 10, sorted and indexed with samtools version 1.16 (Li, et al., 2009; (Danecek, et al., 2021), and conducted single nucleotide polymorphism (SNP) variant calling with bcftools mpileup/call version 1.16 (Danecek, et al., 2021), with maximum depth of 500 and skipping indels, removing bases with base quality less than 30, and reads with mapping quality less than 20. We used the consensus caller, assumed prior heterozygosity of 0.001, and only kept SNPs with probability of data under a null no-SNP model was $p < 0.01$.

We filtered reads with filtering scripts from (Gompert, et al., 2025) and kept biallelic SNPs with ≥ 2x mean depth per individual, missingness per locus no greater than 10%, minimum alternative allele count ≥ 10, BaseQRankSum ≤ 3 and ≥ -3 , MQRankSum ≤ 3 and ≥ -3, ReadPosRankSum ≤ 3 and ≥ -3, and minimum mapping quality ≥ 30. We then filtered to remove SNPs with high coverage (more than 3 standard deviations from the mean) and individuals with mean coverage < 2.5th percentile or > 90th percentile. We converted the VCF output to a genotype likelihood file with a custom script (vcf2gl.pl) and conducted Bayesian genotype estimation using expectation maximization program estpEM version 1.0 (Soria-Carrasco, et al., 2014) with a convergence tolerance of 0.001 and 50 EM iterations. We used glgenest.pl to calculate Bayesian estimates of genotype means from estpEM using the HW allele frequency priors (Soria-Carrasco, et al., 2014).

*Local PCA Clustering*

We clustered windows from our lostruct (Li & Ralph, 2019) analysis within MDS axes if they had the same directionality in z score and two or fewer windows between outlier regions. We then ran a permutation to compute significance based on the longest observed run of outlier windows against a null where the same number of outlier windows is randomly scattered across the genome (n = 1000, p threshold = 0.01), only keeping significant regions. Finally, we filtered to only keep regions with at least 5 consecutive windows. Within mountain, we grouped clusters across MDS axes on the same chromosome if they had any overlap. That region was then assigned the earliest start position and latest end position across windows. Between mountains (Refugio and Hwy154), we grouped inversions if the smallest inversion was at least 2/3 the size of the largest, and at least 80% position overlapping, to match the comparative alignment clustering.

#### Whole-Genome Comparative Alignment

We used the structural variant (SV) calling program SyRI for pairwise SV summaries (Goel, Sun, Jiao , & Schneeberger, 2019). For fused chromosomes in the non-reference genome, we manually fused the chromosomes in the reference to match the orientation. Then, with the output we recalculated the position in the reference coordinate space by splitting and sometimes inverting the manually fused chromosome (https://github.com/dianatataru/Timema-SV-Methods/blob/main/invert_chroms.py).

We used a custom R script (https://github.com/dianatataru/Timema-SV-Methods/blob/main/inversion.R ) for clustering across pairwise comparisons, combining SVs within type (inversion, duplication, translocation) across pairwise comparisons, starting with the largest inversion, if all SVs in the cluster are at least 2/3 the size of the largest inversion in the cluster, and 80% overlapping in reference position with the largest inversion in the cluster. In other words, if SV A and B initially cluster and A is bigger than B, and then we add C, which is smaller than both, to cluster all together both B and C must overlap 80% with A and be at least 2/3 the size of A. We characterized translocations and duplications using the same clustering and filtering and found 1553 translocations and 2328 duplications. These do not include the inverted translocations and duplications.

#### Pangenome Analysis

We created a pangenome graph using 8 haplotype phased genome assemblies in Minigraph-Cactus version 3.0.1 (Hickey, et al., 2024) containing 45,265,393 snarls, or SV bubbles that do not necessarily start and end at the same nodes. We used VG find (Garrison, et al., 2018) to project the pangenome node bounds for inversions back into the linear genome coordinate spaces for the reference genome (GSH2). We used the reference genome coordinate space to identify bounds of an inversion. For some inversions, the start node was not on the reference path (8/51 inversions), so we used the coordinates of a different genome (the other haplotype from the same individual when possible, 5/8 non-reference path inversions). We estimated the size of the inversion by taking the larger of the reference vs. alternate path lengths. We calculated the reference path length as the difference in base pairs between start and end position, and the alternate path length as the number of base pairs in the alternate path.

Using PanTree (Nowbandegani, et al., 2025) to call SVs in the pangenome graph, we detected 26,794,503 total variants; 2,086,235 (7.8%) insertions, 3,204,697 (12%) deletions, 29 duplications, 20,135,058 (75%) SNPs, 1,034,236 (3.9%) multi-nucleotide polymorphisms, and 334,197 (1.2%) replacements (non-SNP, non-MNP).

We also compared a different SV caller, INVPG-annot, which was specifically developed for identifying inversions in pangenomes (Romain, et al., 2025). INVPG-annot only searches for reverse-complement signals, so the logic behind it is simpler than PanTree. We ran it on the clipped pangenome graph, resulting in 209,656 bubbles after filtering and 403 inversion-annotated bubbles. We extracted information from the output VCF file with bcftools version 1.16 (Danecek, et al., 2021), and compared it to the other methods in an R script (inversions.R). This method identified many very small inversions and deeply underestimated both the size and number of inverted regions in the genome (1% of the genome covered), with a hard upper limit of 100 kb (Supplementary Figure 8). Comparisons of this program in simulated versus real data found large reductions in detection from simulated to real, suggesting more limited detection in the program with complex SVs (Romain, et al., 2025). This may be what is happening in our system as well, and we did not include this method in our main analysis for these reasons.

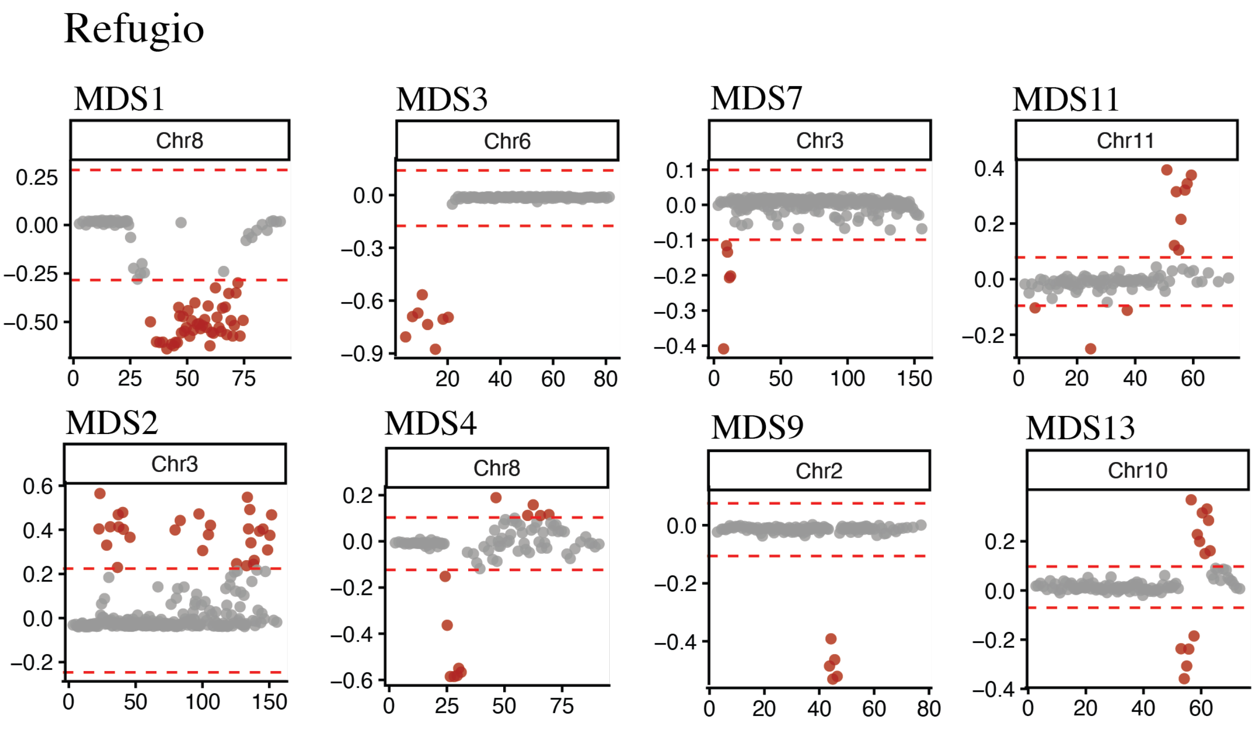

Figure S1. Local PCA MDS outlier runs (putative inversions) for genotyping-by-sequencing data from Refugio.

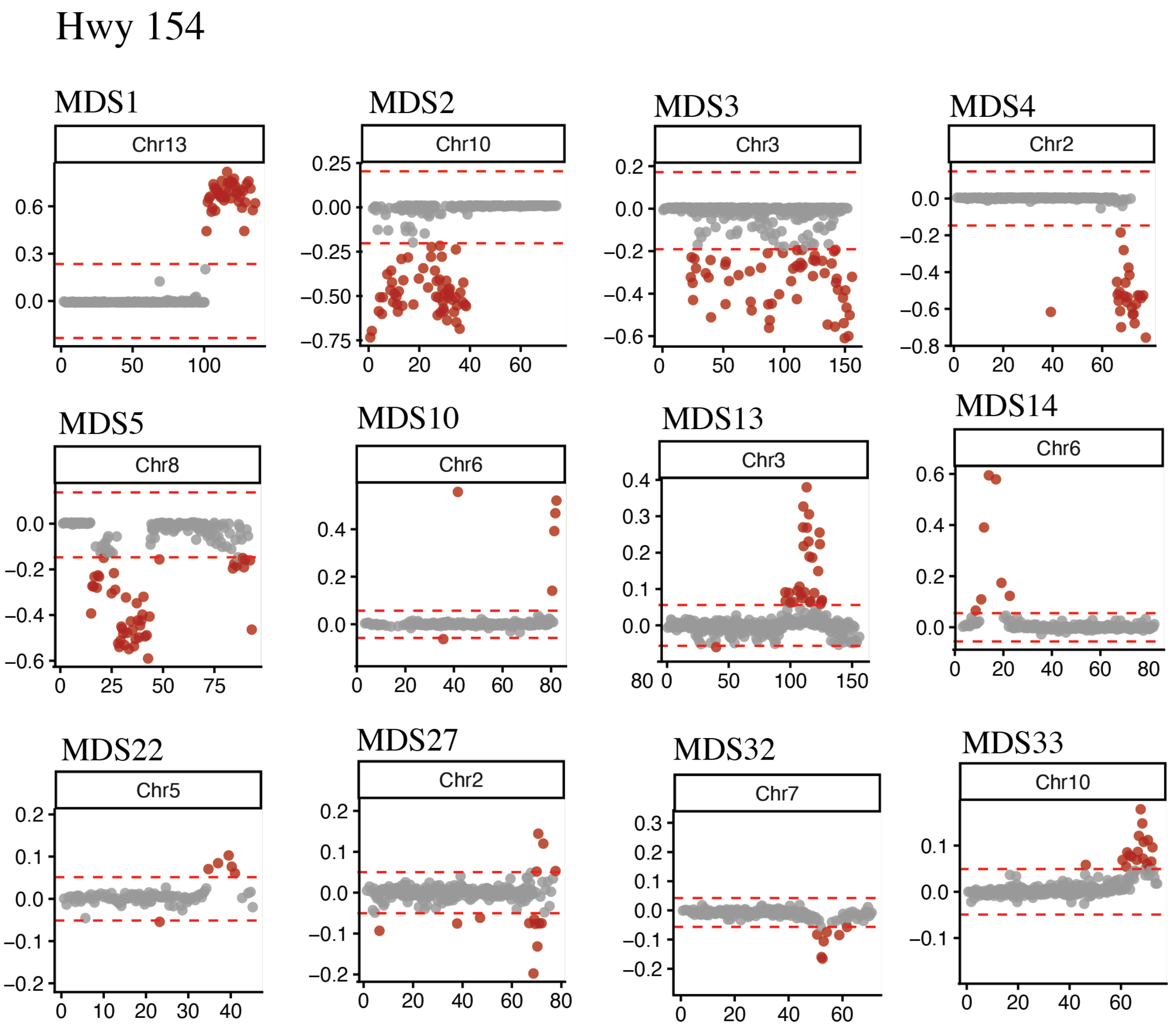


Figure S2. Local PCA MDS outlier runs (putative inversions) for genotyping-by-sequencing data from Hwy 154.

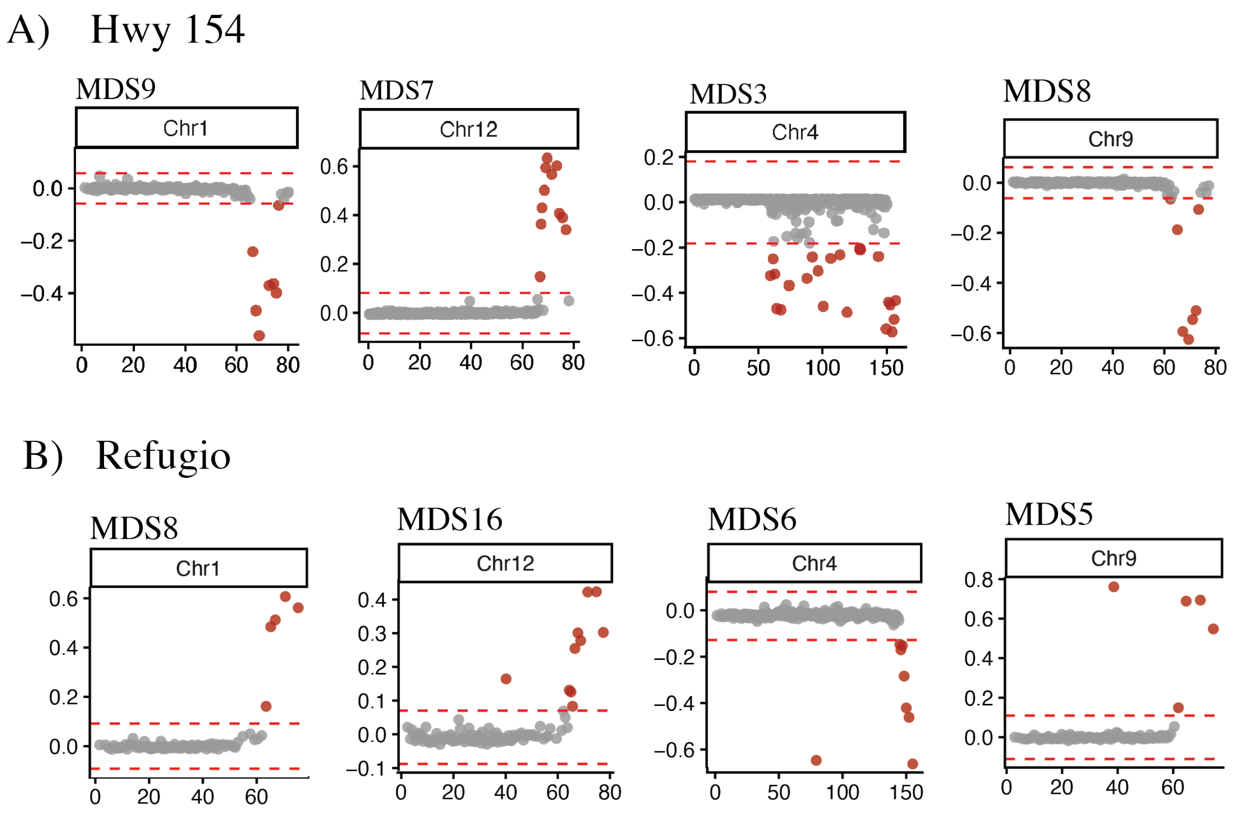


Figure S3. Local PCA MDS outlier runs for genotyping-by-sequencing data for the four shared putative inversions between (A) Hwy 154 and (B) Refugio.

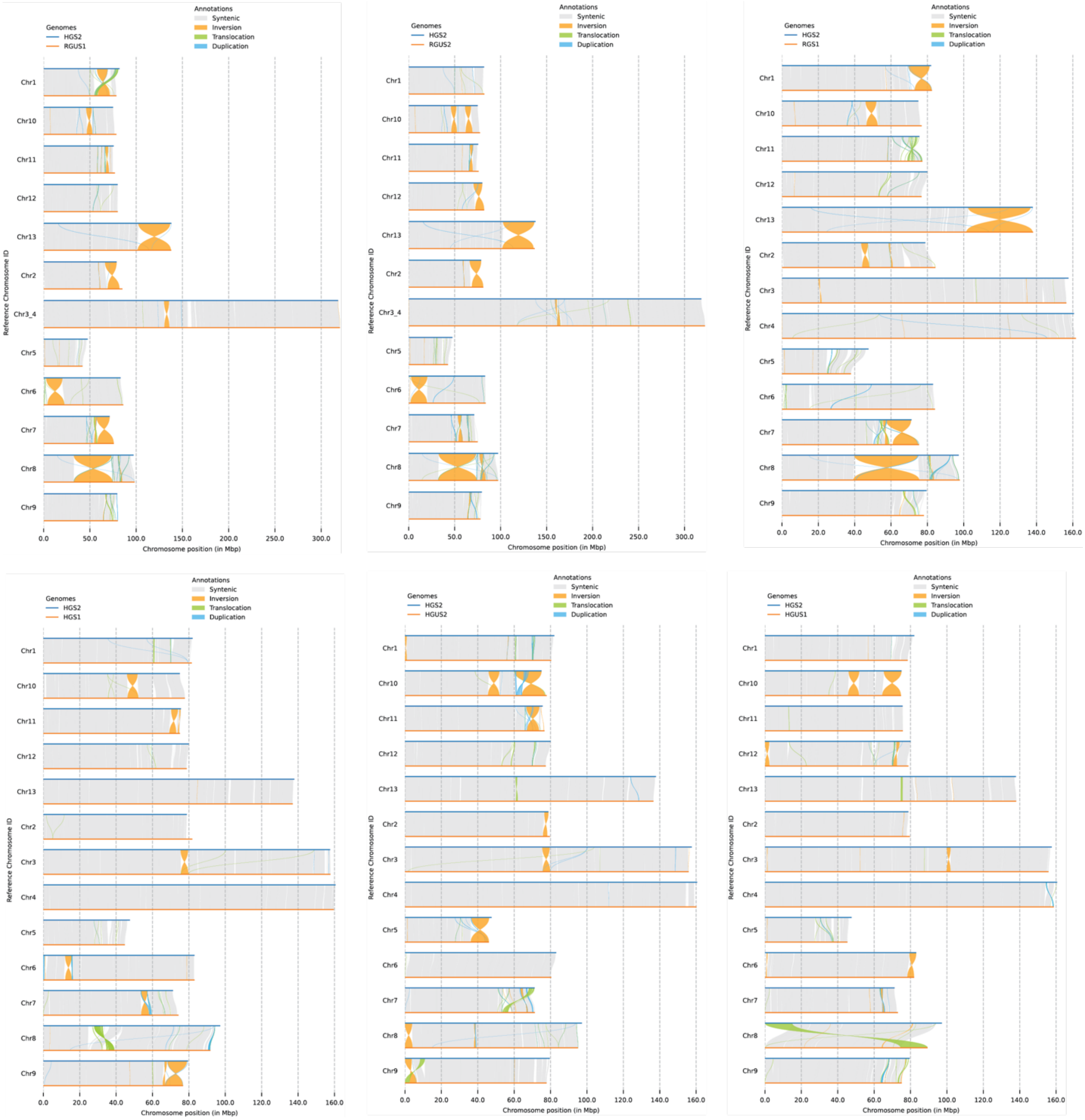


Figure S4. Pairwise comparative alignments between six *de novo* whole-genome assemblies and a reference (GSH2) with structural variant calls across chromosomes from SyRI annotated for inversions (yellow), translocations (green), and duplications (blue).

Chromosome 1

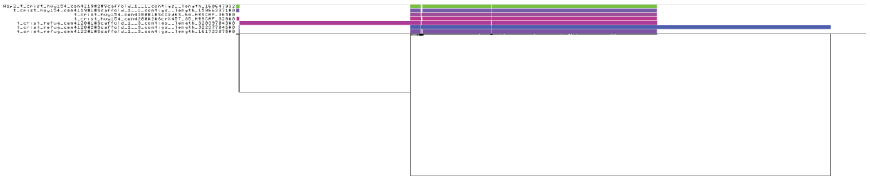

Chromosome 2

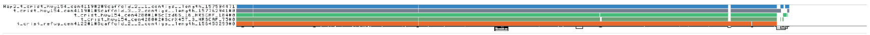

Chromosome 3

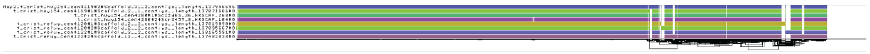

Chromosome 4

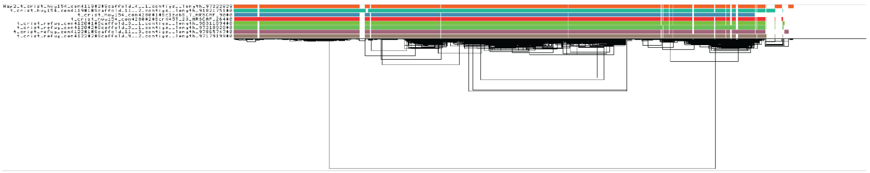

Chromosome 5

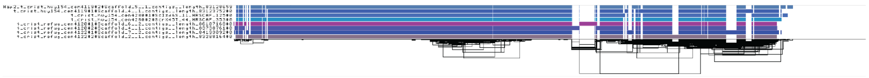

Chromosome 6

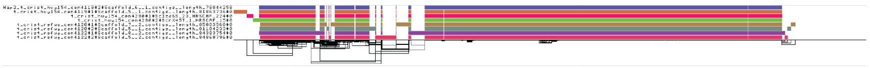

Chromosome 7

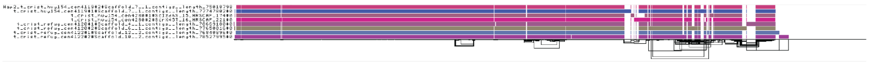

Chromosome 8

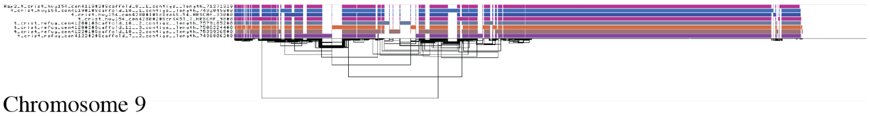

Chromosome 9

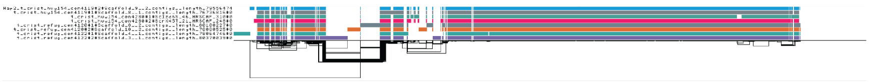

Chromosome 10

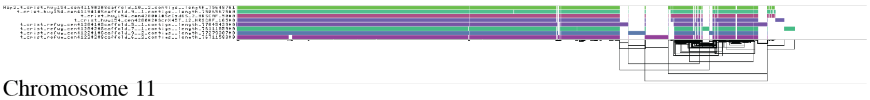

Chromosome 11

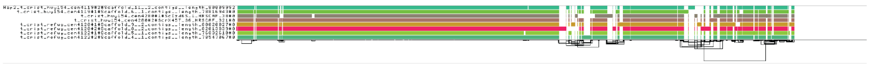

Chromosome 12

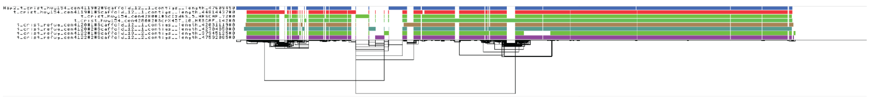

Chromosome 13

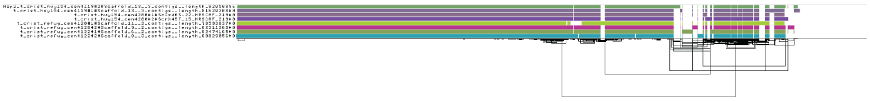

Figure S5. Pangenome graph visualization for all chromosomes (1-13) constructed from the eight genome assemblies visualized in ODGI viz (Optimized Dynamic Genome Graph Implementation visualization; https://github.com/vgteam/odgi). Each bar represents a path (genome) and lines (links) represent graph topology. Genomes are in the same order but are colored differently across chromosomes. Graphs have been sorted using path-guided stochastic gradient descent.

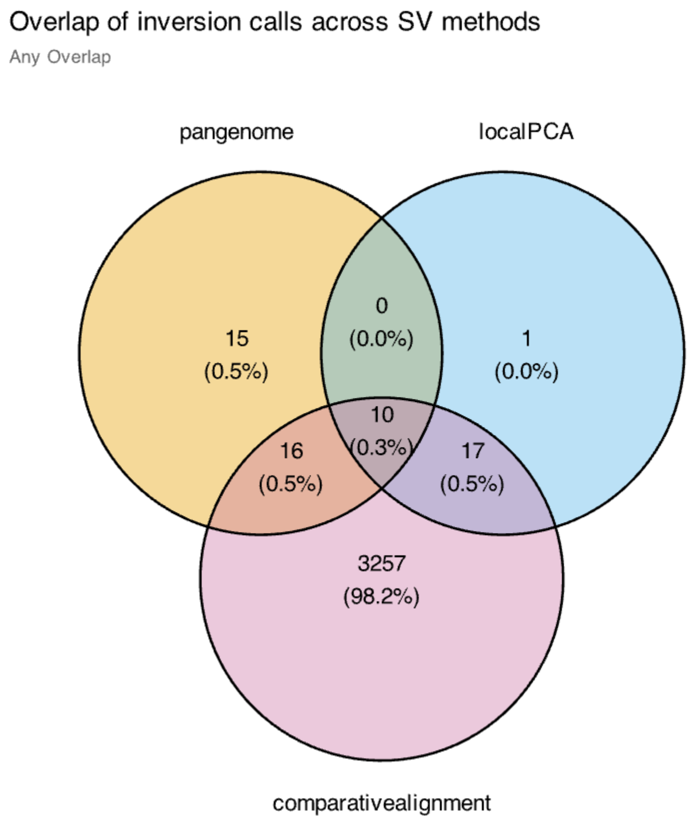


Figure S6. Shared inversions across methods where overlap is defined as any positional overlap.

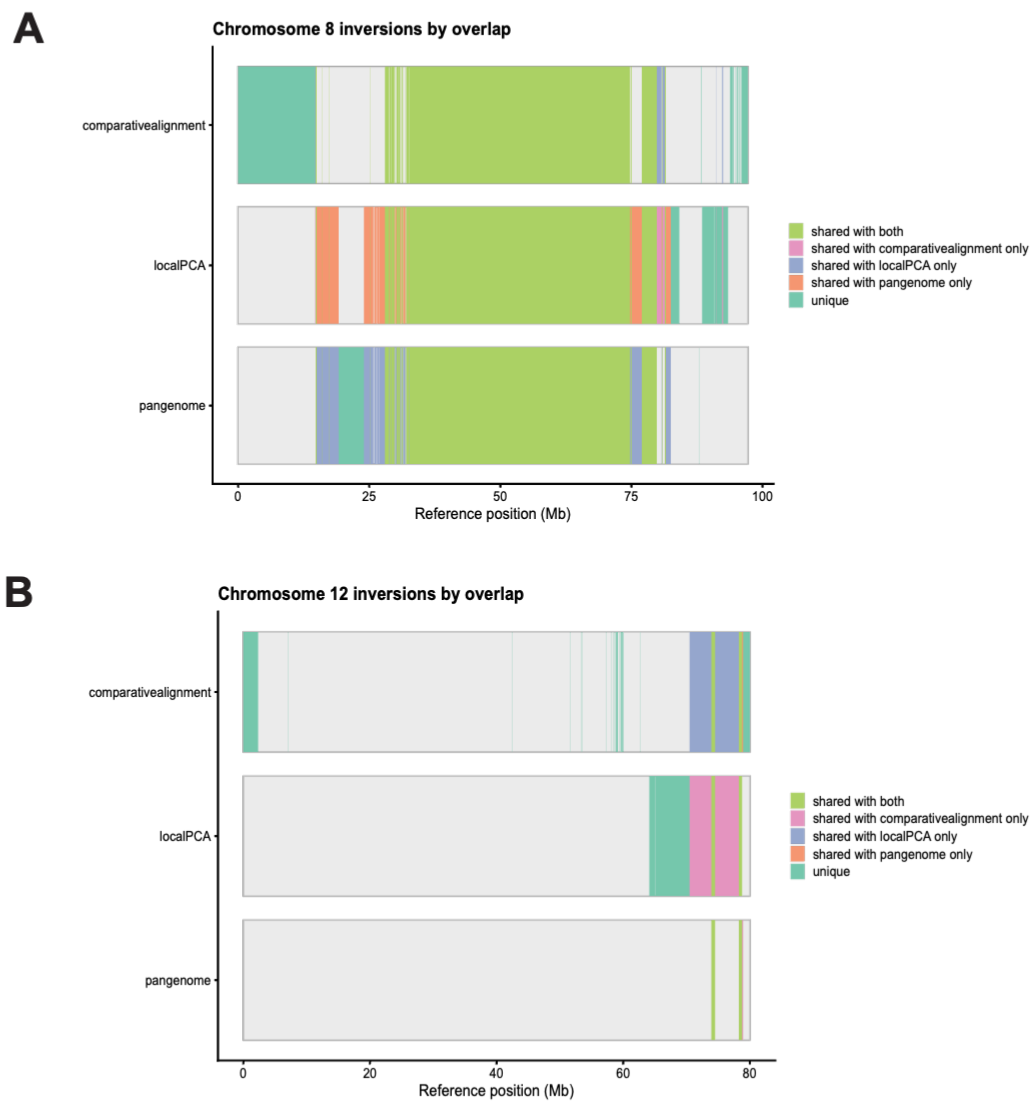


Figure S7. Example chromosome level plots of inversions detected across methods, with inversions colored by whether that inversion location is shared across methods for A) Chromosome 8 and B) Chromosome 12, showing how a large, inverted region can be identified by all three methods, but appear limited in the pangenome, one large block in the local PCA, and many small inversions in the comparative alignment. This is representative of overall patterns across the chromosomes.

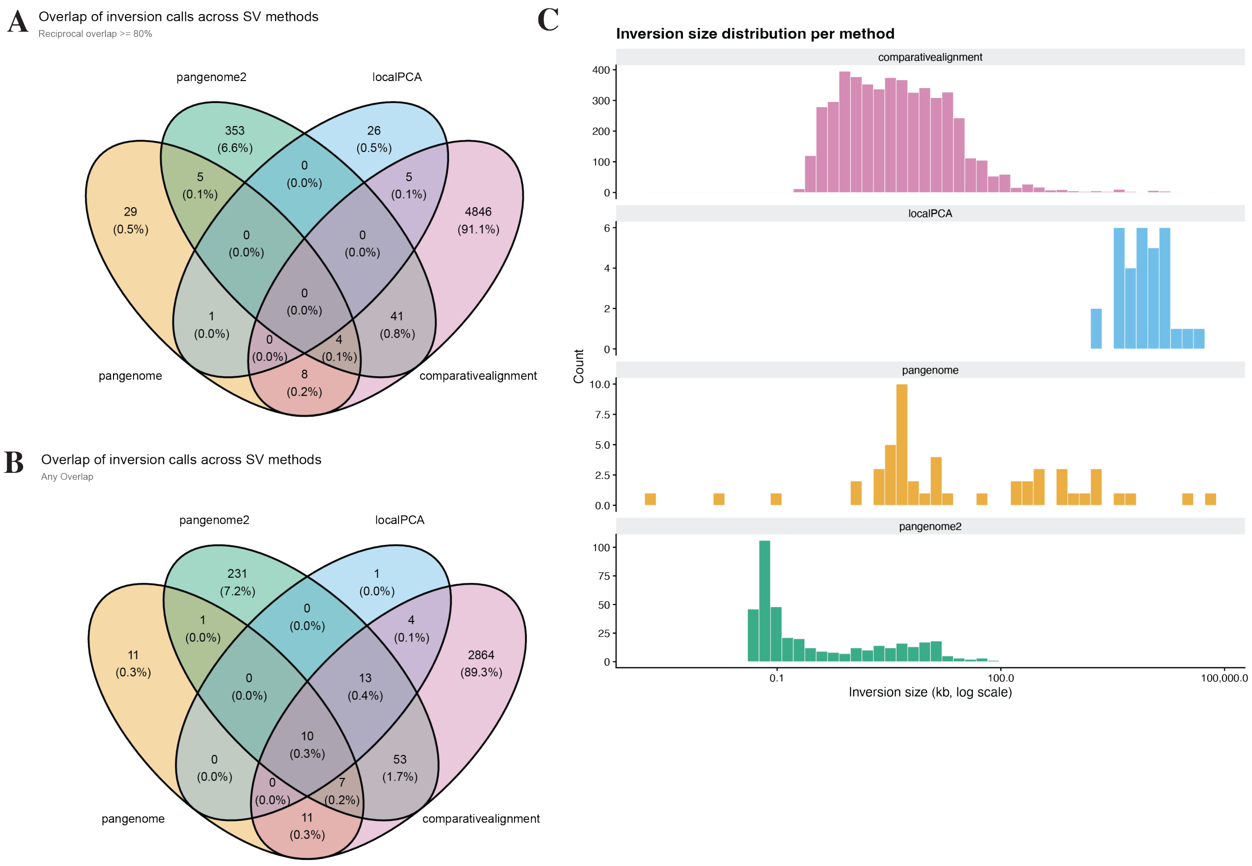


Figure S8. Analyses with the fourth pangenome SV caller, INVPG-annot ("pangenome2" in the figure). Comparison of overlap across methods including an additional SV caller for the pangenome, INVPG-annot (n=403 inversions) where A) overlap is calculated as 80% size and 2/3 position overlap and visualized with a Venn diagram, and B) overlap is calculated by any overlap in position. C) Inversion size distributions (kb, log scale) across all four methods.
